# Rational design of HER2-targeted combination therapies to reverse drug resistance in fibroblast-protected HER2+ breast cancer cells

**DOI:** 10.1101/2024.05.18.594826

**Authors:** Matthew D. Poskus, Jacob McDonald, Matthew Laird, Ruxuan Li, Kyle Norcoss, Ioannis K. Zervantonakis

**Affiliations:** Department of Bioengineering, University of Pittsburgh, Pittsburgh, PA; Hillman Cancer Center, University of Pittsburgh, Pittsburgh, PA; McGowan Institute of Regenerative Medicine, University of Pittsburgh, Pittsburgh, PA

**Keywords:** drug discovery, cancer therapeutics, cell-cell signaling, tumor microenvironment, fibroblasts, systems biology, proteomics

## Abstract

**Introduction:** Fibroblasts, an abundant cell type in the breast tumor microenvironment, interact with cancer cells and orchestrate tumor progression and drug resistance. However, the mechanisms by which fibroblast-derived factors impact drug sensitivity remain poorly understood. Here, we develop rational combination therapies that are informed by proteomic profiling to overcome fibroblast-mediated therapeutic resistance in HER2+ breast cancer cells.

**Methods:** Drug sensitivity to the HER2 kinase inhibitor lapatinib was characterized under conditions of monoculture and exposure to breast fibroblast-conditioned medium. Protein expression was measured using reverse phase protein arrays. Candidate targets for combination therapy were identified using differential expression and multivariate regression modeling. Follow-up experiments were performed to evaluate the effects of HER2 kinase combination therapies in fibroblast-protected cancer cell lines and fibroblasts.

**Results:** Compared to monoculture, fibroblast-conditioned medium increased the expression of plasminogen activator inhibitor-1 (PAI1) and cell cycle regulator polo like kinase 1 (PLK1) in lapatinib-treated breast cancer cells. Combination therapy of lapatinib with inhibitors targeting either PAI1 or PLK1, eliminated fibroblast-protected cancer cells, under both conditions of direct coculture with fibroblasts and protection by fibroblast-conditioned medium. Analysis of publicly available, clinical transcriptomic datasets revealed that HER2-targeted therapy fails to suppress PLK1 expression in stroma-rich HER2+ breast tumors and that high PAI1 gene expression associates with high stroma density. Furthermore, we showed that an epigenetics-directed approach using a bromodomain and extraterminal inhibitor to globally target fibroblast-induced proteomic adaptions in cancer cells, also restored lapatinib sensitivity.

**Conclusions:** Our data-driven framework of proteomic profiling in breast cancer cells identified the proteolytic degradation regulator PAI1 and the cell cycle regulator PLK1 as predictors of fibroblast-mediated treatment resistance. Combination therapies targeting HER2 kinase and these fibroblast-induced signaling adaptations eliminates fibroblast-protected HER2+ breast cancer cells.

## Introduction

The human epidermal growth factor (HER) family plays a critical role in the development and progression of breast cancer [1]. HER2 is overexpressed or amplified in 15-20% of all invasive breast cancer cases and advances in HER2-targeted therapy development have improved survival outcomes [2]. However, residual disease and personalization of HER2-targeted combination therapies remain major challenges in advanced HER2+ breast cancers [2]. Numerous mechanisms involving tumor-intrinsic and tumor-extrinsic factors have been identified that support this drug-resistant state [3, 4]. In the clinic, high abundance of fibroblasts in the HER2+ breast tumor microenvironment has been associated with poor patient outcomes and resistance to multiple therapies [5-9]. Hence, there is a critical need to design effective combination therapies that restore drug sensitivity in these fibroblast-rich breast tumor microenvironments.

Previous studies have demonstrated how individual growth factors activate bypass signaling pathways in cancer cells that limit therapeutic response to HER-family kinase inhibitors [10]. For example, addition of hepatocyte growth factor (HGF) induced resistance to epidermal growth factor inhibition in lung cancer cells, while heregulin induced resistance to HER2 kinase inhibition in breast cancer cells [11]. Another study demonstrated the heterogeneous effects of individual growth factors in blunting therapeutic response to HER2 kinase inhibition depending on cell line molecular subtype (e.g., HGF for the HER2-enriched subtype and heregulin for luminal-HER2 subtype) [12]. Direct coculture of fibroblasts with breast cancer cells has also been linked with reduced sensitivity to HER2-therapy *in vitro* [7, 13-15]. Furthermore, the density [16], origin [17] and matrix-remodeling ability [18] of fibroblasts have been shown to impact breast cancer cell drug sensitivity *in vitro*. Given these heterogeneous responses across breast cancer and fibroblast models, it is critical to examine the response of intracellular signals within cancer cells to the complex fibroblast secretome. In addition, combination therapies to reverse fibroblast-mediated resistance in large panels of HER2+ breast cancer models remain poorly studied and the effects of combination therapies on fibroblast viability have not been investigated.

Here, we systematically evaluate the effects of fibroblast-conditioned medium on the response of a large panel of HER2+ breast cancer cell lines to the FDA-approved HER2 kinase inhibitor lapatinib. Based on these proteomic measurements in fibroblast-protected cancer cells, we design combination therapies and evaluate their ability to restore drug sensitivity. Candidate combination targets were selected using three approaches: (1) a commonly upregulated protein (plasminogen activator inhibitor 1, PAI1) across all cell lines, (2) a protein identified by multivariate data-driven modeling (polio like kinase 1, PLK1) and (3) an epigenetics-directed approach. We examined the effects of these HER2-therapy combinations on both cancer cell and fibroblast viability. Our study reveals a proteomics-informed framework to identify and rationally target fibroblast-mediated mechanisms of drug resistance across multiple HER2+ breast cancer cell lines.

## Materials and Methods

### Cell Culture

Breast cancer cell lines were grown in Roswell Park Memorial Institute (RPMI) 1640 medium supplemented with 10% heat-inactivated fetal bovine serum (HI-FBS) and 1% penicillin (100units/mL)/streptomycin (100ug/mL). Cancer cell lines were engineered to express H2B-GFP (EFM192, HCC1569, MDA-MB-361) or H2B-RFP (BT474, HCC1954, HCC202). AR22 mammary fibroblasts were cultured in Dulbecco’s modified Eagle’s medium (DMEM) supplemented with 10% HI-FBS and 1% penicillin/streptomycin. All cell lines were cultured in a humidified incubator at 5% CO2 and 37C.

### Fibroblast-Conditioned Medium

Conditioned medium was generated by seeding 450000 AR22 fibroblasts in a 15cm plate. After five days, the DMEM medium was aspirated and replaced with RPMI. Conditioned RPMI was harvested four days later and filtered through a 0.2μm filter before storage at -80C. Immediately prior to use in drug-response and proteomic assays conditioned medium was diluted in RPMI to a final concentration of 33% fibroblast-conditioned medium.

### Drug-Response Assay

Cancer cells were seeded in a 96-well black cell culture microplate at 2000 cells/well in either RPMI (monoculture and coculture) or conditioned medium in 100μL. In coculture conditions, an additional 2000 fibroblasts/well were added (1:1 cancer:fibroblast ratio). After 48 hours an additional 100μL of medium was added and plates were treated with lapatinib and inhibitors using a D300 digital dispenser (Tecan, USA). Sytox Deep Red (Invitrogen, USA) was added to media at the time of dosing (final concentration 50nM) to identify dead cells. To quantify cell viability, widefield fluorescence images were acquired after 96 hours of treatment using an ImageXpress Micro confocal in the GFP and Cy5 or RFP and Cy5 channels for H2B-GFP or H2B-RFP cancer cells, respectively. Cell viability was quantified using an Ilastik pixel classifier trained to identify the area of viable cells (GFP+/Cy5-or RFP+/Cy5-) in each image. Area under the curve (AUC) was computed from drug response of seven lapatinib doses (0.003-3μM) normalized to DMSO cell numbers. To assess fibroblast viability, plates were fixed in 4% paraformaldehyde and subsequently incubated with 2μg/mL Hoechst 33342 for 15 minutes to stain nuclei. Fluorescence images in the DAPI channel were acquired with a Nikon Ti2 microscope and the number of fibroblast nuclei were quantified using Cellprofiler.

### Analysis of Reverse-Phase Protein Array Data

Cancer cells were seeded at 0.2M cells/well in a 6-well plate in either RPMI or conditioned medium. After 48 hours, cells were treated with DMSO or 0.1μM lapatinib for 48 hours. Plates were washed twice with ice-cold PBS prior to protein lysis. Reverse-phase protein analysis was performed following previous protocols [19]. Partial least squares regression was performed using the pls library in Rstudio. Protein abundance of lapatinib-downregulated proteins in cancer cells cultured in fibroblast-conditioned medium under lapatinib treatment (BT474, EFM192, HCC1569, HCC202, MDA-MB-361) was z-scored and was used as the predictor variable and subsequently regressed against AUC (response variable).

### Bioinformatic Analysis

Data from the METABRIC study [20] was acquired from cBioPortal. HER2+ breast cancer patients were filtered based on clinical annotation and deconvolution was performed in R with the package immunedeconv to determine the stromal score of each sample. PAI1 gene expression levels were correlated to the stroma score using Pearson’s correlation coefficient. The gene expression matrix and clinical annotations of the GSE130788 dataset were acquired from GEO with the R package GEOquery. The pre-therapy gene expression data was used to calculate the stroma scores (estimate) and to separate patients in high or low stroma groups based on the median value. PLK1 gene expression levels pre-/post-HER2-therapy changes were compared between the high and low stroma groups.

## Results

### Fibroblast-derived paracrine factors heterogeneously modify lapatinib sensitivity and proteomic responses in a panel of HER2+ breast cancer cell lines

We used a panel of nine HER2+ breast cancer cell lines and evaluated their sensitivity to the FDA-approved HER2 kinase inhibitor lapatinib under conditions of monoculture (cancer cells only) or exposure to fibroblast-conditioned medium. All cancer cell lines exhibited a dose-dependent reduction in viable cell numbers with increasing concentration of lapatinib (**Fig 1A**). However, the baseline sensitivity was variable among these cell lines with the minimum lapatinib dose that reduced cell numbers ranging between 3nM to 100nM (**Fig 1A**). Lapatinib response in the presence of fibroblast-conditioned medium remained unaffected in four out of the nine cancer cell lines that were termed as “fibroblast-insensitive” (**Fig 1B**, AU565, HCC1419, HCC1954, UAC812). The remaining five HER2+ breast cancer cells exhibited higher cell viability at multiple lapatinib doses and were termed as “fibroblast-protected”. The lapatinib dose at which fibroblast-conditioned medium increased cell viability ranged between 10nM-1μM for MDA361, 30nM-1μM for BT474, 100nM-3μM for EFM192 or HCC202 and 300nM-3μM for HCC1569. We also computed the area under the curve (AUC) metric, with higher AUC values indicating higher cell viability (**Fig 1B**). We found that fibroblast-insensitive and fibroblast-protected HER2+ breast cancer cells exhibited a similar range of AUC values under monoculture (fibroblast-insensitive: 2.94-3.84 vs. fibroblast-protected: 2.49-3.99).

**Figure 1.**
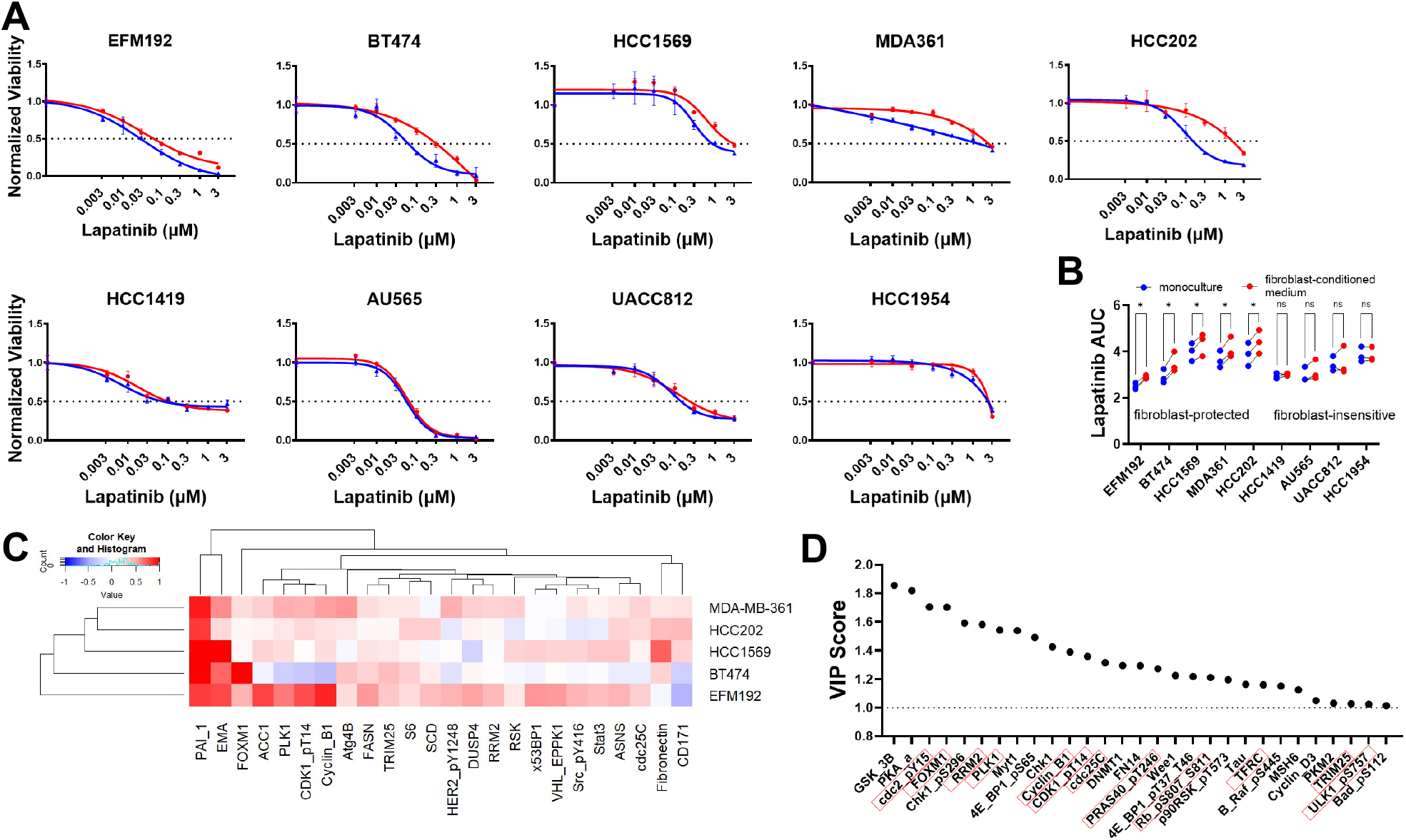
Heterogeneous effects of fibroblast-conditioned medium on HER2+ breast cancer cell sensitivity and proteomic response to lapatinib. **(A)** Dose-response of nine HER2+ breast cancer cell lines to lapatinib in monoculture (blue) or cultured in fibroblast-conditioned medium (red). Viable cells were measured 96hrs after treatment with lapatinib. Error bars represent replicate wells and results are representative of three biological replicates. **(B)** Area under the curve (AUC) values for the dose-response curves in panel (A). Dots represent paired biological replicates and cancer cell lines are classified as “fibroblast-protected” vs. “fibroblast-insensitive”. Paired t-test, *: p<0.05. (**C**) Heatmap of proteins that are upregulated by fibroblast-conditioned medium compared to monoculture in cancer cells treated with lapatinib. Out of 385 proteins profiled, 24 were upregulated in at least two cancer cell lines. Values represent the log2-normalized ratio of protein expression for fibroblast-conditioned medium normalized to monoculture. Both conditions were treated with 100nM lapatinib. **(D)** Ranked variable importance (VIP) scores for proteins that predict lapatinib AUC using a partial least squares regression model (proteins that were identified by both the differential expression analysis and VIP>1 are shown with a red box outline).

Next, using reverse phase protein arrays we identified proteins in the five fibroblast-protected cancer models that were upregulated by fibroblast-conditioned medium under conditions of lapatinib treatment. Out of the 383 proteins profiled, a total of 24 proteins were upregulated by 1.2-fold in at least two breast cancer models (**Fig 1C** and **Table S1** shows all 69 proteins upregulated in at least one cancer cell line). Plasminogen activator inhibitor-1 (PAI1) was the only protein upregulated in all five cancer cell lines and there are no studies linking PAI1 with HER2 therapy resistance. Epithelial membrane antigen (EMA) was upregulated in four cancer cell lines and epithelial-to-mesenchymal transitions have been linked with HER2 therapy resistance [21, 22]. Fibronectin was upregulated in three cancer cell lines and has been previously shown to mediate resistance to HER2 kinase inhibition [23]. In addition, acetyl-coa-carboxylase 1 (ACC1) was upregulated in three cancer cell lines and regulates fatty acid synthesis that has been linked with resistance to HER2-targeted therapies [24]. Finally, another 20 proteins were upregulated in at least two cancer cell lines, with multiple proteins previously associated with resistance to HER2 targeted therapy including forkhead box protein M1 [25], ribonucleotide reductase M2 [26], signal transducer and activator of transcription 3 [7], ribosomal protein S6 [27], and fatty acid [28].

To further investigate proteomic predictors of poor response to lapatinib in the context of fibroblast-protection, we employed partial least squares regression (PLSR). PLSR accounts for co-variation between proteins (x-variable) that can be linked with drug response (y-variable) and represents a data-driven approach to uncover candidate targets for combination therapies [11, 29-31]. In this multivariate regression analysis, 29 out of 98 proteins (limited out of 383 proteins by selecting those downregulated by lapatinib) exhibited a higher-than-average contribution [32] in predicting lapatinib sensitivity (**Fig 1D**, variable importance score > 1). Consistent with our differential protein expression analysis, 13 out of these 29 proteins had higher expression in the fibroblast-conditioned medium condition compared to monoculture. Among these overlapping proteins multiple were related to control of cell cycle, including polo like kinase 1 (PLK1), cell division cycle 2 (cdc2), and cyclin dependent kinase 1 (CDK1), supporting previous studies on the role of cell cycle in HER2 therapy resistance [33]. Collectively, our results demonstrate that fibroblast-derived factors modulate response to lapatinib in a subset of HER2+ breast cancer cell lines and that proteomic profiling of the fibroblast-protected state uncovers previously known targets of HER2 therapy resistance.

### PAI1 expression in cancer cells is commonly upregulated by fibroblast-conditioned medium and PAI1 pharmacologic blockade improves response to HER2 kinase therapy

The uniform pattern of PAI1 upregulation induced by fibroblast-conditioned medium led us to evaluate whether fibroblast-protected breast cancer cells are sensitive to PAI1 blockade. First, we evaluated the efficacy of single-agent activity of tiplaxtinin, a selective small molecule inhibitor of PAI1 [34], on breast cancer cells cultured in the presence of fibroblast-conditioned medium. We found that tiplaxtinin reduced the number of viable cancer cells in a dose-dependent manner (**Fig 2A**). Reduction of cell viability at an intermediate tiplaxtinin dose of 20μM varied between 15% (BT474, least sensitive) and 49% (EFM192, most sensitive. This effect did not exhibit any association with cancer cell response to HER2-therapy since both cancer cells with low (EFM192) or high (HCC1569) lapatinib AUC values exhibited similar sensitivity to tiplaxtinin.

**Figure 2.**
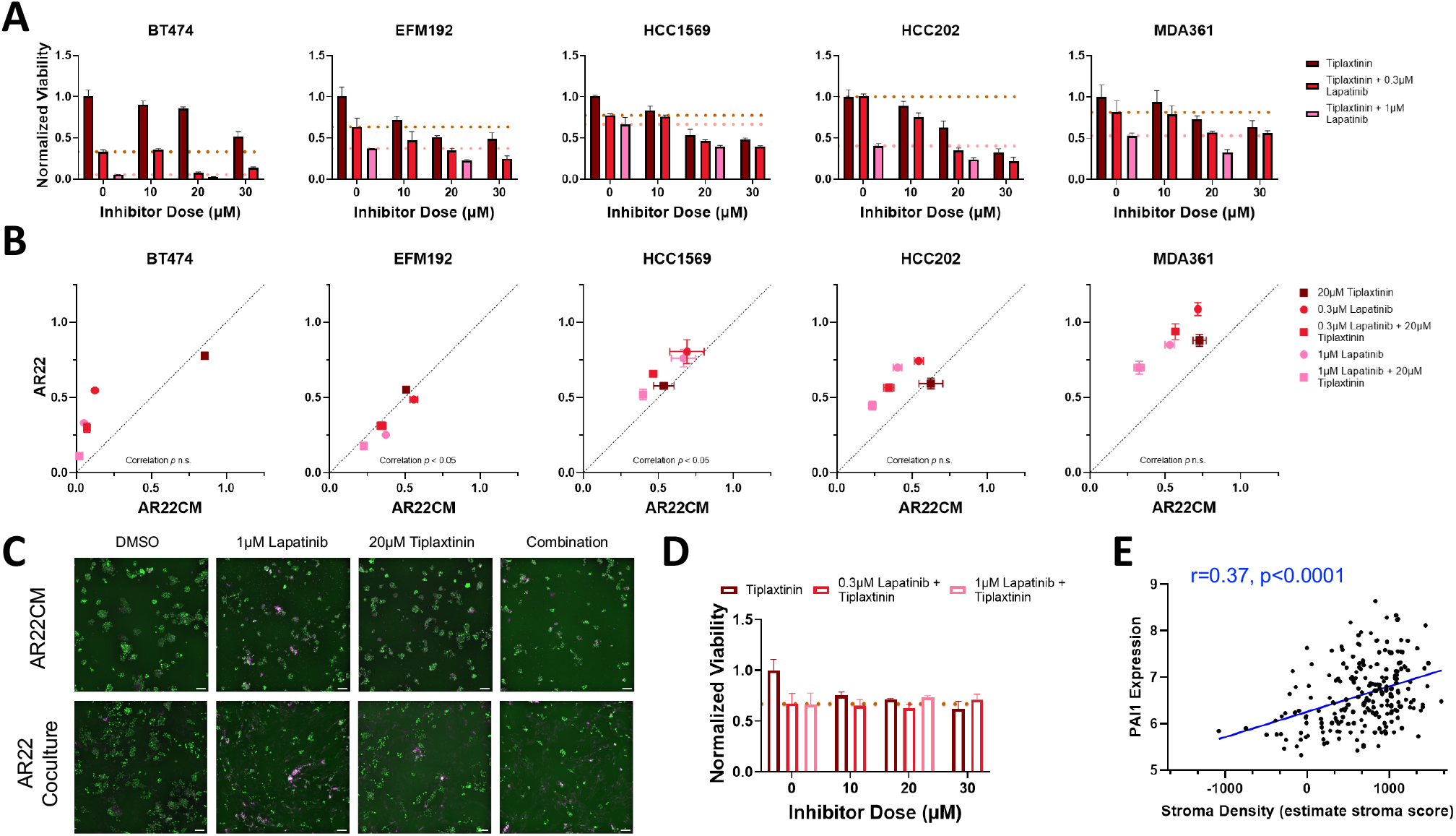
PAI1 inhibition increases efficacy of lapatinib in fibroblast-protected HER2+ breast cancer cells. **(A)** Cancer cell viability in response to the single-agent PAI inhibitor tiplaxtinin (dark red) and in combination with a low (0.3μM, red) or high (1μM, pink) lapatinib dose. Dashed lines represent the cell viability values for each lapatinib dose as monotherapy. Results are shown for each fibroblast-protected cancer cell line (error bars are standard error of the mean for replicate wells) and are representative for three biological replicates. **(B)** Relationship between cancer cell viability under conditions of direct coculture with fibroblasts (y-axis, AR22) and culture in fibroblast-conditioned medium (x-axis, AR22CM) for two lapatinib doses and one tiplaxtinin dose as single agents or in combination. **(C)** Representative images of EFM192 breast cancer cells (labeled with a nuclear H2B-GFP marker (green) and stained with dead cell marker (magenta) for the conditions in panel (B) at the indicated doses. Scale bar is 100μm. **(D)** Viability of fibroblasts with increasing dose of tiplaxtinin. Results are shown for each fibroblast-protected cancer cell line (error bars are standard error of the mean for replicate wells) and are representative for three biological replicates. **(E)** PAI1 gene expression levels correlate with stroma density in HER2+ breast cancer patients (METABRIC dataset, n=224 tumors, pearson correlation r=0.37, p<0.0001).

We next assessed cancer cell response to tiplaxtinin in combination with either 300nM or 1μM lapatinib. The addition of 20μM tiplaxtinin with a low lapatinib dose (300nM) potently reduced cancer cell survival compared to lapatinib monotherapy in four out of the five breast cancer cell lines (all except MDA361). The HER2 kinase/PAI1 inhibition combination was most lethal to BT474 cells, resulting in reduction of viable cells by 93% in both cell lines compared to 67% by lapatinib as a single agent. In contrast, the addition of tiplaxtinin reduced the viability of MDA361 cells by 46% versus 29% reduction by lapatinib monotherapy at the low lapatinib dose. A higher lapatinib dose (1μM) combined with tiplaxtinin (20μM) reduced cancer cell survival compared to lapatinib alone for all cell lines. Consistent with their response to tiplaxtinin plus 300nM lapatinib, BT474 was the most sensitive cell lines to this dose and exhibited a decrease of 98% in viable cells respectively versus a 95% reduction by lapatinib alone. EFM192 and HCC202 exhibited intermediate sensitivity to this combination and their viability was reduced by 78% and 77% compared to a reduction of 63% and 60% respectively from 1μM lapatinib alone. HCC1569 cells were least sensitive to this dose and the addition of 20μM tiplaxtinin to 1μM lapatinib improved the response from 33% reduction in cell viability in lapatinib monotherapy to 60%.

We further evaluated the potential of PAI1-targeted therapy on eliminating fibroblast-protected cancer cells under conditions of direct coculture with fibroblasts. Consistent with the conditioned medium results, cancer cell viability was lowest for the combined tiplaxtinin and lapatinib treatment compared to either monotherapy (**Fig 2B-C**). The number of surviving cancer cells after treatment in direct coculture and conditioned medium correlated (pearson correlation coefficient r between 0.61-0.90 for all cell lines); however, HCC1569 and MDA361 cells were less sensitive to all treatments in coculture compared to conditioned medium. Despite this, the most potent combination of 20μM tiplaxtinin plus 1μM lapatinib reduced HCC1569 and MDA361 viability by 49% and 30% in direct coculture. As a critical control comparison, we also examined the effects of PAI1 and HER2 kinase combination therapy on fibroblast viability and found lower reduction in cell viability compared to breast cancer cells (**Fig 2D:** 27% reduction in fibroblast viability in 20μM tiplaxtin combined with 1μM lapatinib compared to a range of 30% to 89% reduction in cancer cells cocultured with fibroblasts).

In addition, we examined the association of PAI1 and stroma scores using the publicly available METABRIC dataset that included transcriptomic data from n=224 HER2+ breast cancer patients. We found that tumors with high PAI1 gene expression also exhibited a high stroma score, indicative of a fibroblast-rich microenvironment (**Fig 2E**, pearson correlation coefficient r=0.37). Taken together, these findings demonstrate that blockade of PAI1 improves response to HER2 kinase inhibition across multiple HER2+ breast cancer cell lines under both conditions of direct coculture and exposure to fibroblast-conditioned medium.

### Fibroblast-secreted factors sustain PLK1 protein expression in cancer cells treated with lapatinib and inhibiting PLK1 restores sensitivity to lapatinib

Next, we examined the therapeutic potential of targeting the cell cycle regulator PLK1 that was identified both by differential expression analysis and multivariate partial-least squares regression as a predictor of poor lapatinib response in fibroblast-protected cancer cells. Out of all cell cycle regulators identified, PLK1 was selected because it is an actionable target and has not been previously examined in the context of fibroblast-mediated therapy resistance. We measured the response of the five HER2+ breast cancer cell lines grown in fibroblast-conditioned medium to the ATP-competitive PLK1 inhibitor GSK461364 [35] alone or in combination with lapatinib. All cancer cell lines exhibited a dose-dependent reduction in the number of viable cells after four days of PLK1 pharmacologic blockade. Response to 10nM GSK461364 monotherapy was heterogenous among the five cell lines: sensitivity ranged from a maximum of 66% reduction in the number of viable cells in the most sensitive cell line HCC202 to a modest reduction of 12% and 15% at this dose in MDA361 and HCC1569 cells, respectively (**Fig 3A**). At the highest dose (100nM) cell viability was inhibited in all cell lines with a range of 62% to 81%.

**Figure 3.**
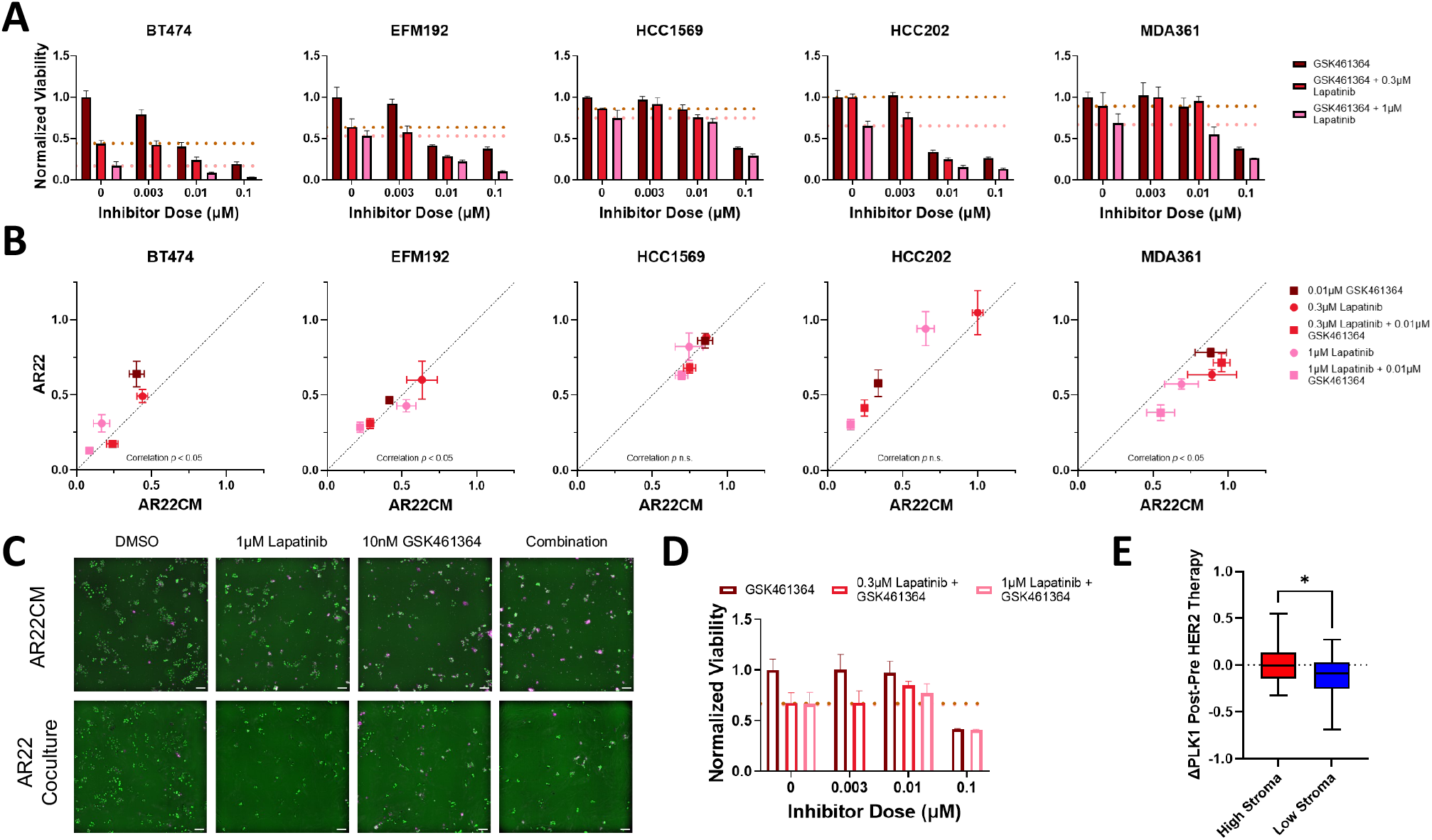
Targeting PLK1 restores lapatinib sensitivity in fibroblast-protected HER2+ breast cancer cells and PLK1 represents a clinically-relevant vulnerability in stroma-rich HER2+ breast tumors. **(A)** Cancer cell viability in response to the single-agent PLK1 inhibitor GSK461364 (dark red) and in combination with a low (0.3μM, red) or high (1μM, pink) lapatinib dose. Dashed lines represent the cell viability values for each lapatinib dose as monotherapy. Results are shown for each fibroblast-protected cancer cell line (error bars are standard error of the mean for replicate wells) and are representative for three biological replicates. **(B)** Relationship between cancer cell viability under conditions of direct coculture with fibroblasts (y-axis, AR22) and culture in fibroblast-conditioned medium (x-axis, AR22CM) for two lapatinib doses and one GSK461364 dose as single agents or in combination. **(C)** Representative images of EFM192 breast cancer cells (labeled with a nuclear H2B-GFP marker (green) and stained with dead cell marker (magenta) for the conditions in panel (B) at the indicated doses. **(D)** Viability of fibroblasts with increasing dose of GSK461364. Results are shown for each fibroblast-protected cancer cell line (error bars are standard error of the mean for replicate wells) and are representative for three biological replicates. Scale bar is 100μm. **(E)** HER2-therapy does not effectively reduce PLK1 gene expression levels in HER2+ patients with high stroma (red) compared to low stroma (blue). The y-axis indicates the difference in PLK1 gene expression levels post HER2-therapy minus pre HER2-therapy. Data from GSE130788 dataset (n=89 paired pre/post-therapy tumors) and high vs. low stroma groups separated by the mean expression estimate stroma score.

We next assessed the combined effects of PLK1 inhibition plus either 300nM or 1μM lapatinib. A dose of 10nM GSK461364 plus 300nM lapatinib greatly reduced the number of viable cells only in three cell lines: BT474, EFM192, and HCC202 compared to lapatinib monotherapy. This combination was equally potent in BT474 and HCC202, resulting in a 76% reduction in cell numbers despite differential sensitivity to lapatinib alone; 300nM lapatinib monotherapy did not reduce the viability of HCC202 but induced a 56% reduction in BT474. Conversely, dual HER2 kinase/PLK1 blockade modestly increased the viability reduction of HCC1569 cells and did not alter response of MDA361 cells compared to lapatinib alone. Combining 10nM GSK461364 with a higher lapatinib dose (1μM) significantly reduced viable cells compared to lapatinib or GSK461364 monotherapy in all five cell lines. BT474 was most sensitive to this combination and treatment reduced the number of tumor cells at endpoint by 92% compared to 84% reduction by lapatinib alone. HCC1569 tumor cells were least sensitive to this dose and their viability was inhibited by 30% versus 25% by lapatinib monotherapy.

We next treated all five cancer cell lines cultured with fibroblasts to assess how direct coculture with fibroblasts affects cancer cell response to lapatinib plus PLK1 inhibition. The response of all five cell lines directly cultured with fibroblasts generally correlated with response to cancer cells cultured in fibroblast-conditioned medium (**Fig. 3B-C**, pearson correlation coefficient r between 0.86 and 0.96 for HCC1569 and HCC202, respectively). Treatment with 10nM GSK461364 and 1μM lapatinib induced a response ranging between 37% reduction in HCC1569 cells and 87% reduction in BT474 in direct coculture versus 31% and 92% for HCC1569 and BT474 in fibroblast-conditioned medium. EFM192 and HCC1569 exhibited similar drug sensitivity in fibroblast-conditioned medium and direct coculture, whereas BT474 and HCC202 were less sensitive to monotherapy and combination therapy when directly cocultured with fibroblasts compared to conditioned medium. MDA361 cells were marginally but consistently less sensitive to lapatinib monotherapy and combination therapy in conditioned medium versus direct coculture. We also evaluated the dose-response of dual HER2 kinase/PLK1 inhibition on fibroblast viability. GSK461364 minimally impacted the number of fibroblasts at endpoint at doses less than 100nM and the addition of 1μM lapatinib with 10nM GSK461364 reduced viability by 23% (**Fig. 3D**).

The relationship between PLK1 sustained expression following HER2 therapy and fibroblasts was further examined using a publicly available transcriptomic dataset from the TRIO-US B07 clinical trial of HER2+ breast cancer patients treated with HER2-targeted therapies (n=89 pre-/post-therapy tumors with transcriptomic data [36]). We found that in fibroblast-rich tumors PLK1 gene expression levels were similar pre- and post-therapy compared to a reduction of PLK1 expression induced by therapy in tumors with low fibroblast density (**Fig 3E**). Thus, PLK1 represents a clinically relevant target in HER2+ breast tumors and its inhibition sensitizes multiple fibroblast-protected HER2+ breast cancer cell lines to lapatinib.

### Combining epigenetic-based BET bromodomain inhibition with lapatinib eliminates fibroblast-protected HER2+ breast cancer cells

Given that each fibroblast-protected breast cancer cell line exhibited heterogeneous patterns of proteomic changes in response to fibroblast-conditioned medium (**Fig 1C** and **Table S1**), we next evaluated the potential of an epigenetic-based drug combination to globally mitigate these heterogeneous responses. Based on previous studies of lapatinib-induced tumor cell autonomous kinome reprogramming in HER2+ breast cancer cell lines [37], we selected to evaluate the effects of JQ1, a bromodomain and extraterminal domain (BET) inhibitor [38], on fibroblast-protected cancer cells. We first evaluated the response of all five cancer cell lines cultured in fibroblast-conditioned medium exposed to 100nM-1μM JQ1 for four days. We found that JQ1 reduced cell viability compared to baseline in a dose-dependent manner (**Fig 4A)**. Sensitivity was heterogeneous among cell lines and JQ1 reduced viability between 26-32% compared to baseline for the least sensitive cell line (BT474) and between 45-71% for the most sensitive cell line (EFM192) at the lowest and highest doses, respectively.

**Figure 4.**
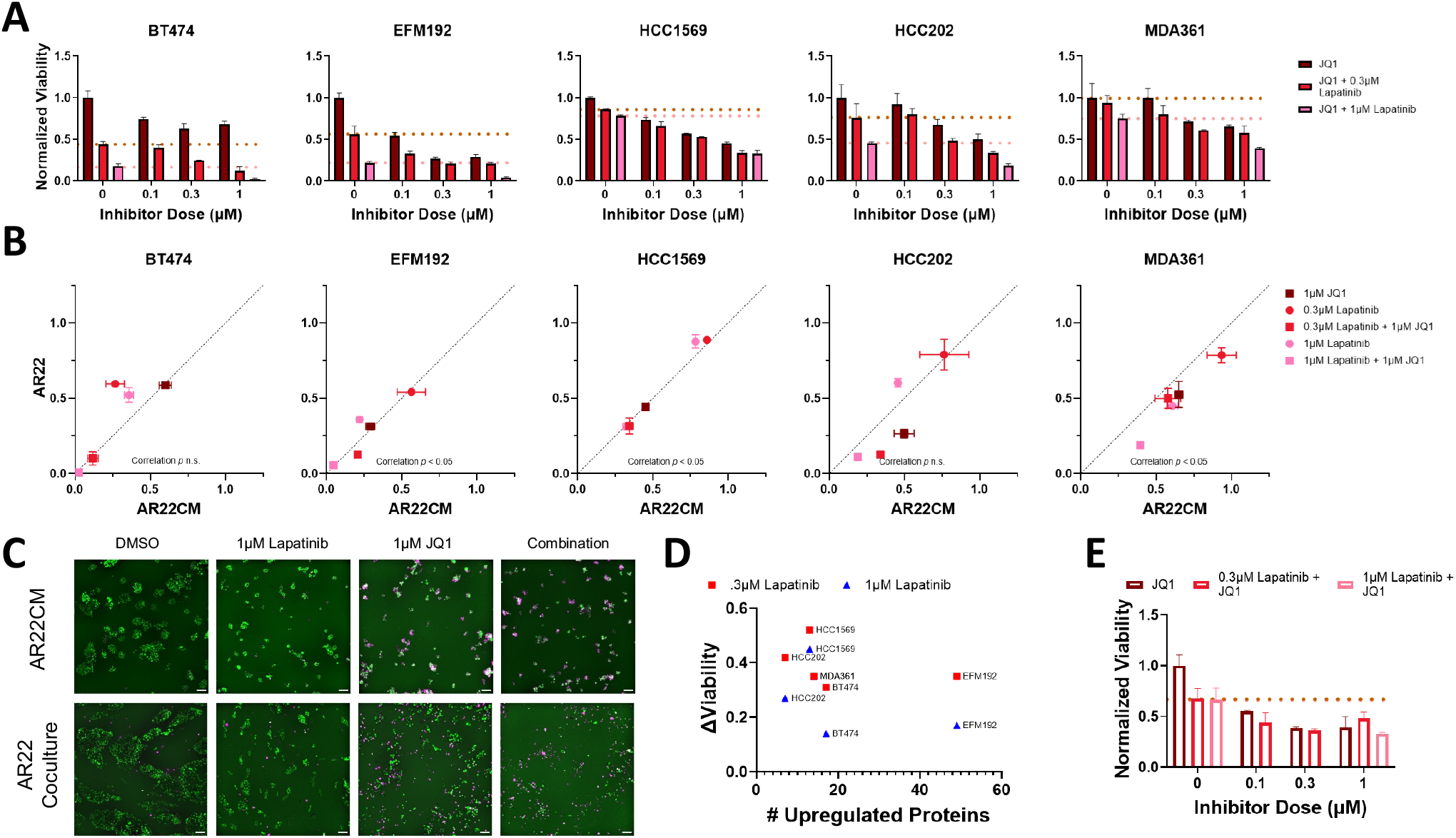
Combination therapy with the epigenetic inhibitor JQ1 and lapatinib eliminates fibroblast-protected HER2+ breast cancer cells. **(A)** Cancer cell viability in response to the BET inhibitor JQ1 as a single agent (dark red) and in combination with a low (0.3μM, red) or high (1μM, pink) lapatinib dose. Dashed lines represent the cell viability values for each lapatinib dose as monotherapy. Results are shown for each fibroblast-protected cancer cell line (error bars are standard error of the mean for replicate wells) and are representative for three biological replicates. **(B)** Relationship between cancer cell viability under conditions of direct coculture with fibroblasts (y-axis, AR22) and culture in fibroblast-conditioned medium (x-axis, AR22CM) for two lapatinib doses and one JQ1 dose as single agents or in combination. **(C)** Representative images of EFM192 breast cancer cells (labeled with a nuclear H2B-GFP marker (green) and stained with dead cell marker (magenta) for the conditions in panel (B) at the indicated doses. Scale bar is 100μm. **(D)** Relationship between number of upregulated proteins and sensitization by JQ1 in all fibroblast-protected breast cancer cell lines. **(E)** Viability of fibroblasts with increasing dose of JQ1. Results are shown for each fibroblast-protected cancer cell line (error bars are standard error of the mean for replicate wells) and are representative for three biological replicates.

To evaluate efficacy of BET bromodomain/HER2 kinase inhibition on cancer cells in the presence of fibroblast-secreted factors, we treated cells with 1μM JQ1 plus either 300nM or 1μM lapatinib and measured the reduction in viable cells (**Fig 4B**). Combination of JQ1 plus low lapatinib dose (300nM) greatly reduced the number of surviving tumor cells for all cell lines compared to lapatinib monotherapy. BT474, EFM192, and HCC202 tumor cells were highly sensitive to this combination and survival of these cells was greatly reduced compared to both JQ1 and lapatinib monotherapy. BT474 was the most sensitive of these three cell lines and combination therapy reduced viable cancer cells by 88% compared to 56% by lapatinib alone. MDA361 were comparatively less sensitive than all other cell lines, but the combination was significantly more effective than lapatinib monotherapy and resulted in a 42% reduction in cell numbers compared to 7% reduction with 300nM lapatinib alone. Pairing 1μM JQ1 with the higher lapatinib dose (1μM) also decreased the number of surviving tumor cells compared to lapatinib alone. These conditions nearly completely eradicated EFM192 and BT474, reducing viable cells by 95% (versus 78% in lapatinib monotherapy) and 97% (versus 82% in lapatinib monotherapy), respectively. The number of surviving HCC1569 and HCC202 cells was also potently reduced by 67% and 81% at this dose. MDA361 cells were the least sensitive to these conditions and the JQ1/lapatinib combination decreased the number of cells by 61% versus 25% by lapatinib alone.

Dual BET bromodomain/HER2 kinase inhibition was also effective in reducing tumor cell numbers in direct coculture conditions and correlated with response in fibroblast-conditioned medium conditions (**Fig 4B-C**, pearson correlation coefficient r between 0.71-0.99). Consistent with the fibroblast-conditioned medium results, BT474 and EFM192 cells cocultured with fibroblasts were highly sensitive to the most potent combination of 1μM JQ1 and 1μM lapatinib treatment and their viability was reduced by 99% and 94%, respectively. Furthermore, we examined whether the extent of fibroblast-induced proteomic changes in cancer cells was associated with the ability of JQ1 to potential the effects of lapatinib. We found that cancer cell lines with a similar reduction in cell viability (e.g., BT474 and EFM192) exhibited different number of fibroblast-induced upregulated proteins (**Fig 4D**). Finally, we evaluated the impact of JQ1 alone and in combination with lapatinib on fibroblast viability. JQ1 monotherapy significantly reduced fibroblast numbers between 45-61% in a dose-dependent manner. Combination treatment reduced fibroblast viability by a maximum of 67% at 1μM JQ1 and 1μM lapatinib, which was less potent than the effects in BT474 and EFM192, but comparable to effects in HCC1569, HCC202, MDA361 (**Fig 4E)**. Overall, these findings support the efficacy of JQ1 plus lapatinib combination in reducing cancer cell numbers in the presence of either fibroblasts or fibroblast-conditioned medium.

## Discussion

A better understanding of cancer cell survival mechanisms in fibroblast-rich breast tumor microenvironments is necessary to develop new therapies that improve patient outcomes in advanced HER2+ breast cancer. Profiling the drug-resistant state in a complex tumor ecosystem allows for discovery of exploitable vulnerabilities in cancer cells [39]. In this study, we utilized a panel of HER2+ breast cancer cell lines to define how fibroblast-conditioned medium impacts cancer cell proteomic responses following treatment with lapatinib. We selected to target either proteins that were upregulated by fibroblast-derived paracrine factors or an epigenetic regulator. We evaluated the effects of HER2-kinase combination therapies with inhibitors of these targets in both cancer-fibroblast cocultures and cancer cells exposed to fibroblast-conditioned medium. Our integrated proteomic and drug response profiling enabled the development of rational combination therapies to restore drug sensitivity in fibroblast-protected HER2+ breast cancer cells.

Fibroblasts represent an abundant stromal cell type in the breast tumor microenvironment [5]. Previous investigations have shown that direct coculture of breast cancer cells with fibroblasts limits sensitivity to multiple anticancer therapies, including chemotherapy [16-18] and kinase-targeted therapies [7]. In addition to evaluating direct coculture and paracrine factors, previous studies focused on the effects of individual fibroblast-derived signals, such as heuregulin [11, 12, 40]. Our results on fibroblasts protecting only a subset of breast cancer cell lines from HER2-targeted therapy are supported by a previous study that demonstrated the heterogeneous effects of extracellular ligands in the response of HER2+ breast cancer cells to lapatinib [12]. Furthermore, transcriptomic analysis in a single HER2+ breast cancer cell line cocultured with fibroblasts has previously shown activation of the TGF-β and WNT signaling pathways [7]. However, no studies have investigated the effects of the complex fibroblast secretome on intracellular cancer cell proteomic responses and developed proteomics-informed combination therapies to reverse resistance to anti-HER2 therapy.

In our studies, fibroblast-conditioned medium consistently upregulated PAI1 across all five fibroblast-protected breast cancer cell lines, however PAI1 has not been previously studied in the context of HER2-therapy resistance. PAI1 is a member of the serine protease inhibitor (Serpin) family (encoded by *SERPINE1*) and is upregulated in breast tumors compared to benign breast tissue [41]. Furthermore, PAI1 together with urokinase plasminogen activator (uPA), represents a clinically validated biomarker and high PAI1/uPA levels predict a higher rate of recurrence in lymph node-negative disease treated with chemotherapy [42]. Importantly, high PAI1 gene expression levels have been associated with poor outcomes in HER2+ breast cancer patients [43]. Our results of PAI1 contributing to the survival of fibroblast-protected cancer cells in the context of HER2 therapy resistance agree with previous reports in esophageal and pancreatic carcinomas that demonstrated PAI1-targeted therapy restored cancer cell sensitivity to chemotherapy [44, 45]. Furthermore, although no previous study has examined the role of PAI1 in breast cancer-fibroblast crosstalk, adipocytes have been shown to upregulate PAI1 expression in cancer cells [46]. In addition, the pro-tumorigenic role of PAI1 is supported by the finding that genetic silencing of PAI1 in both cancer cells and the host tissue reduces tumor growth *in vivo* [47]. Collectively, these findings highlight PAI1 as a stroma-regulated therapeutic target that associates with aggressive disease in HER2+ breast cancer.

As an additional approach to exploit therapeutic vulnerabilities in fibroblast-protected breast cancer cells, we explored targeting PLK1 that predicted lapatinib response in cancer cells exposed to fibroblast-conditioned medium. The kinase PLK1 regulates cell division and is under clinical investigation for multiple tumor types (ongoing trials NCT05768932, NCT05358379 and completed trials reviewed [48]). However, PLK1-based HER2 combination therapies have not been investigated in the context of fibroblast-mediated resistance. Preclinical studies have shown that genetic and pharmacologic blockade of PLK1 in HER2+ breast cancer cells restored sensitivity to the anti-HER2 antibody-drug conjugate TDM1 [49]. Another study demonstrated that PLK1 inhibition synergizes with taxane-based chemotherapy *in vitro* and *in vivo* [50]. In the clinic, high PLK1 levels have been associated with resistance to cyclin-dependent kinase 4/6-targeted therapy in luminal breast cancer [51], and to estrogen-targeted therapy [52]. Taken together, these studies position PLK1 as a promising combination therapy partner to enhance treatment response to multiple anticancer treatments in breast cancer, including chemotherapy, estrogen- and HER2-targeted therapy.

## Conclusion

We present an integrated approach of proteomic and drug response profiling to rationally design HER2-targeted combination therapies that eliminate fibroblast-protected HER2+ breast cancer cells. We demonstrate three approaches (PAI1-targeted, PLK1-targeted, and epigenetics-targeted) to restore HER2 therapy sensitivity in a panel of fibroblast-protected HER2+ breast cancer cells. Our findings that the proteolytic regulator PAI1 and cell cycle kinase PLK1 drive fibroblast-induced therapy resistance are also supported by transcriptomic analyses of HER2+ breast tumors in the clinic.

## Acknowledgements

This work was supported by the National Institute s of Health (R00 CA222554 to I.K.Z and T32 EB001026 to M.D.P.), the Department of Bioengineering and Swanson School of Engineering at the University of Pittsburgh. We thank Dr. Gordon Mills, Dr. Yiling Lu and the MD Anderson Functional Proteomics Reverse Phase Protein Array Facility (supported by P30CA016672 and R50CA221675) for providing the proteomic data.

## Statements and Declarations

## Conflict of interest

The authors declare no competing interests.

## Notes

### Competing Interest Statement

The authors have declared no competing interest.

